# *Drosophila small ovary* encodes a zinc-finger repressor required for ovarian differentiation

**DOI:** 10.1101/354746

**Authors:** Leif Benner, Elias A. Castro, Cale Whitworth, Koen J.T. Venken, Haiwang Yang, Brian Oliver, Kevin R. Cook, Dorothy A. Lerit

**Affiliations:** Section of Developmental Genomics, Laboratory of Cellular and Developmental Biology, National Institute of Diabetes and Digestive and Kidney Diseases, National Institutes of Health, Bethesda MD 20892.; Department of Biology, Johns Hopkins University, Baltimore MD 21218.; Department of Cell Biology, Emory University School of Medicine, Atlanta GA 30322.; Department of Biology, Indiana University, Bloomington, Indiana 47405.; Verna and Marrs McLean Department of Biochemistry and Molecular Biology, McNair Medical Institute at the Robert and Janice McNair Foundation, Dan L. Duncan Cancer Center, Center for Drug Discovery, Department of Pharmacology and Chemical Biology, Baylor College of Medicine, Houston, TX, 77030, USA

**Keywords:** chromatin, differentiation, gene expression, HP1a, oogenesis, position-effect variegation

## Abstract

Repression is essential for coordinated cell type-specific gene regulation and controlling the expression of transposons. In the *Drosophila* ovary, stem cell regeneration and differentiation requires controlled gene expression, with derepression leading to tissue degeneration and ovarian tumors. Likewise, the ovary is acutely sensitive to deleterious consequences of transposon derepression. The *small ovary* (*sov*) locus was identified in a female sterile screen, and mutants show dramatic ovarian morphogenesis defects. We mapped the locus to the uncharacterized gene *CG14438*, which encodes a zinc-finger protein that colocalizes with the essential Heterochromatin Protein 1 (HP1a). We demonstrate that Sov functions to repress inappropriate cell signaling, silence transposons, and suppress position-effect variegation in the eye, suggesting a central role in heterochromatin stabilization.

## Summary statement

*Small ovary* is required for heterochromatin stabilization to repress the expression of transposons and ectopic signaling pathways in the developing ovary.

## Introduction

Multicellular organisms rely on stem cells. A combination of extrinsic and intrinsic signals function to maintain the balance between stem cell self-renewal and differentiation needed for tissue homeostasis of stem cell development. The *Drosophila* adult ovary is a well-studied model for development (Fuller and Spradling 2007). It is organized into ˜16 ovarioles, which are assembly lines of progressively maturing egg chambers. At the anterior tip of each ovariole is a structure called the germarium, harboring 2–3 germline stem cells (GSCs), which sustain egg production throughout the life of the animal. Surrounding the GSCs and their cystoblast daughters is a stem cell niche, a collection of nonproliferating somatic cells composed of terminal filament and cap cells. Like the GSCs, somatic stem cells divide to produce new stem cells and escort cells, which encase the germline. Each cystoblast is fated to differentiate and will undergo four rounds of cell division with incomplete cytokinesis to produce a germline cyst interconnected by the fusome, a branched cytoskeletal structure. Germline cysts are surrounded by two escort cells. Toward the posterior of the germarium, a monolayer of somatic follicle cells replaces the escort cells to form an egg chamber.

Egg chamber development requires the careful coordination of distinct germline and somatic cell populations through signaling within a complex microenvironment. For example, extrinsic transducers and signals including Jak/Stat and the BMP homolog, Decapentapalegic (Dpp), are required to maintain the proliferative potential of GSCs (Bausek 2013; Gilboa 2015; S. Chen, Wang, and Xie 2011). Oriented divisions displace the primary cystoblast cell away from the stem cell signals and permit the expression of differentiation factors, such as *bag of marbles* (*bam*). The importance of Bam is revealed in *bam* loss-of-function mutants, which have tumorous germaria filled with undifferentiated germline cells containing dot spectrosomes, the dot-fusome structure characteristic of GSCs and primary cystoblasts as a result of complete cytokinesis (Lin and Spradling 1995; D. McKearin and Ohlstein 1995). Thus, germline differentiation requires the repression and activation of cell signaling pathways.

Dynamic changes in chromatin landscapes within germline and somatic cells are critical for oogenesis and contribute to the regulated gene expression required for tissue homeostasis (Barton et al. 2016; X. Li et al. 2017; Börner et al. 2016; Peng et al. 2016; Soshnev et al. 2013; McConnell, Dixon, and Calvi 2012). Nucleosomes generally repress transcription by competing for DNA binding with transcription factors (Lorch and Kornberg 2017; Kouzarides 2007; Jenuwein and Allis 2001). The reinforcement of repression depends largely on histone modifications. For example, the activities of both the H3K9 methyltransferase SETDB1, encoded by *eggless* (*egg*) (Wang et al. 2011; Clough et al. 2007; Clough, Tedeschi, and Hazelrigg 2014), and the H3K4 demethylase encoded by *lysine-specific demethylase 1* (Di Stefano et al. 2007; Rudolph et al. 2007; Eliazer, Shalaby, and Buszczak 2011; Eliazer et al. 2014) are required in the somatic cells of the germarium for GSC maintenance, normal patterns of differentiation, and germline development. Genomewide profiling hints at a progression from open chromatin in stem cells to a more closed state during differentiation (T. Chen and Dent 2014). Disrupting this progressive repression in the ovary is predicted to contribute to stem cell overproliferation and defective oogenesis.

There are other roles for regulated chromatin states in oogenesis. The propensity of transposons to mobilize in gonads is countered by host responses, such as induced heterochromatin formation, to repress transposon activity. The proximity of condensed heterochromatin also dampens the activity of genes and transgenes (Elgin and Reuter 2013).

Originally described over 40 years ago (J. D. Mohler 1977), the *small ovary* (*sov*) locus is associated with a range of mutant phenotypes including disorganized ovarioles, egg chamber fusions, undifferentiated tumors, and ovarian degeneration (Wayne et al. 1995). In the present work, we define the molecular identify and function of Sov. We demonstrate *sov* encodes an unusually long C_2_H_2_ zinc-finger (ZnF) nuclear protein. Previous work suggested Sov may complex with the conserved Heterochromatin-Protein 1a (HP1a) (Alekseyenko et al. 2014), encoded by *Su(var)205* and critical for heterochromatin formation (James and Elgin 1986; Eissenberg et al. 1990; Clark and Elgin 1992). RNA-seq analysis showed that *sov* activity is required in the ovary to repress the expression of a large number of genes and transposons. Additionally, Sov functions as a dominant suppressor of position-effect variegation (PEV) in the eye, similar to HP1a (Elgin and Reuter 2013). Moreover, Sov and HP1a colocalize in the nucleus. These data indicate that Sov is a novel repressor of gene expression involved in heterochromatization.

## Results

### small ovary is CG14438

To refine previous mapping of *sov*, we used complementation of female sterile and lethal *sov* mutations with preexisting and custom-generated deficiencies, duplications, and transposon insertions (J. D. Mohler 1977; D. Mohler and Carroll 1984; Wayne et al. 1995). Our results (Fig 1A, B) suggested that either the protein-coding *CG14438* gene or the intronic, noncoding *CR43496* gene is *sov* since the locus mapped to the noncomplementing deletion *Df(1)sov*. Rescue of female sterility and/or lethality of *sov^2^*, *sov^ML150^*, *sov^EA42^* and *Df(1)sov* with *Dp(1;3)sn^13a1^* (molecularly undefined, not shown), *Dp(1;3)DC486*, and *PBac{GFP-sov}* confirmed our mapping. We could not replicate separability of *sov^EA42^* sterility and lethality with *Dp(1;3)sn^13a1^* (Wayne et al. 1995). Female sterile and lethal alleles of *sov* map identically.

**Figure 1.**
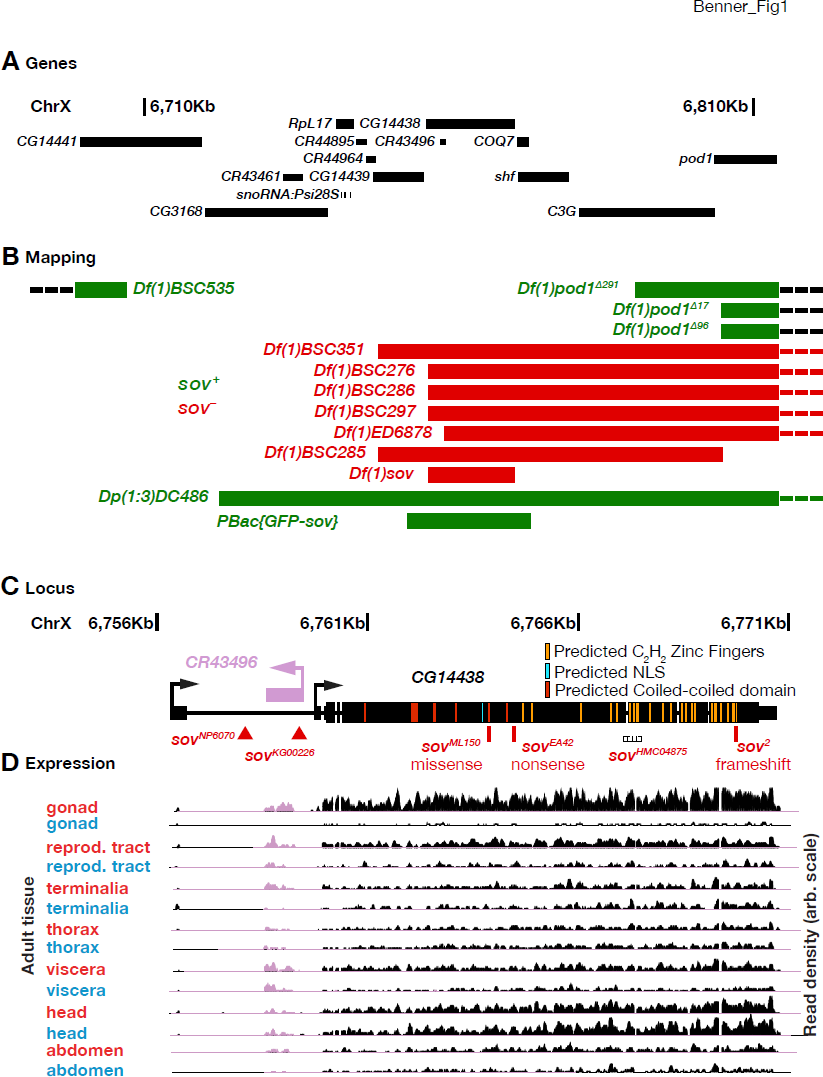
***sov*** **is** ***CG14438***. (A) Genes of the genomic interval X:6710000–6810000 (Gramates et al. 2017). (B) Deficiency (*Df*) and Duplication (*Dp*) mapping. Noncomplementing (*sov*^−^, red) and complementing (*sov^+^*, green) alleles and rearrangements are shown. (C) Schematic of the *CG14438* (black) and *CR43496* (purple) genes. Transcription start (bent arrows), introns (thin lines) non-coding regions (medium lines) and coding regions (thick lines) are shown. Transposon insertions (triangles), point mutations (red lines), the region targeted by the shRNAi transgene (base-paired), and Sov protein features are shown. (D) RNA expression tracks by tissue type from female (red) or male (blue) adults.

We performed genome sequencing to determine if the lesions in *sov* alleles occur in *CG14438* or *CR43496* (Table S1). While *CR43496* contained three polymorphisms relative to the reference genome, we identified the same polymorphisms in all *sov* mutant and control lines. Thus, *CR43496* is unlikely to be *sov.* In contrast, we found disruptive mutations in *CG14438* (Fig 1C), which encodes an unusually long, 3,313-residue protein with 21 C_2_H_2_ ZnFs and multiple nuclear localization motifs and coiled-coil regions (Fig 1C). We identified a nonsense mutation (G to A at position 6,764,462) in the *CG14438* open reading frame of *sov^EA42^* which is predicted to truncate the *sov* protein before the ZnF domains. We found a frameshift insertion (T at position 6,769,742) located towards the end of *CG14438* in *sov^2^* that encodes 30 novel residues followed by a stop codon within the C-terminal ZnF and removes the terminally predicted NLS (Fig 1C). We found a missense mutation (C to G at position 6,763,888) in *sov^ML150^* that results in a glutamine to glutamate substitution within a predicted coiled-coil domain. While this is a conservative substitution and glutamate residues are common in coiled-coil domains, glutamine to glutamate substitutions are especially disruptive in the coiled-coil region of the sigma transcription factor (Hsieh, Tintut, and Gralla 1994). We conclude that *CG14438* encodes *sov*.

### sov is highly expressed in the ovary

To determine where *sov* is expressed and if it encodes multiple isoforms, we analyzed its expression in adult tissues by RNA-seq. We noted that *sov* is broadly expressed as a single mRNA isoform, with highest expression in ovaries (Fig 1D). The modENCODE (Graveley et al. 2011; J. B. Brown et al. 2014) and FlyAtlas (Robinson et al. 2013; Leader et al. 2018) reference sets show a similar enrichment in the ovary and early embryos. The enrichment of *sov* in the ovary is consistent with its reported oogenesis phenotypes.

To determine where Sov is expressed in the ovary, we generated a GFP-tagged transgene (GFP-Sov) sufficient to rescue ovary degeneration in *sov* mutants. Examination of the distribution of GFP-Sov in developing egg chambers (Fig 2A) revealed Sov localization in several cell types. By pairing localization analysis of Sov with antibodies recognizing Vasa (Vas) to label the germline, we observed particularly striking nuclear localization of Sov surrounded by perinuclear Vas in the germline cells within region 1 of the germarium, with highest levels evident within the GSCs (dashed circles; Fig 2B). Using Traffic jam (Tj) to label the soma highlighted nuclear enrichment of Sov in the Tj-positive somatic and follicle stem cells (dashed ovals; Fig 2C). These data indicate that Sov is enriched in the germline and somatic cells of the ovary, including both stem cell populations.

**Figure 2.**
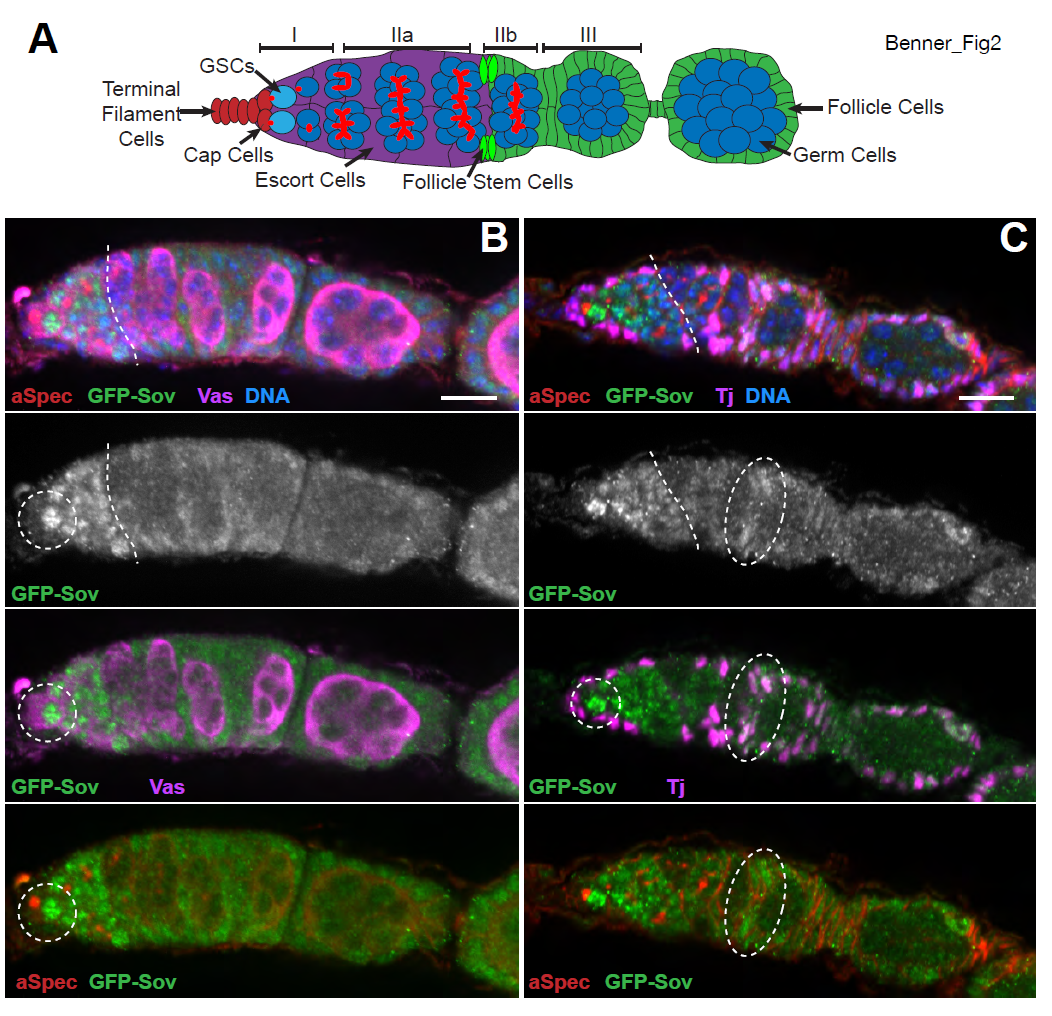
**Sov germarium expression.** (A) Cartoon showing germarium regions I-III and young egg chamber with cell types labeled. (B,C) 1d posteclosion, visualized for GFP-Sov (green), anti-αSpectrin (red). Images show single optical sections. (B) Sov contrasted with anti-Vas germline (magenta), or (C) -Tj somatic staining (magenta). GFP-Sov expression regions of interest (see text) are shown (dashed lines). Bars: 10μm.

### Sov is required in the soma and germline for oogenesis

To examine *sov* function in the germarium, we compared a strong allelic combination, *sov^EA42^*/*sov^2^,* to cell-type specific knockdown of *sov* using a UAS short hairpin *sov* RNAi construct (P{TRiP.HMC04875}; hereafter, *sov^RNAi^*). The germaria of heterozygous *sov* females are wild type (Fig 3A), but *sov* mutants often show an ovarian tumor phenotype, with greater than the expected number of germ cells with dot spectrosomes (Fig 3B), indicating that germ cells undergo complete cytokinesis. Ovarian tumors were observed in ˜40% of *sov* mutants (N=12/31 *sov^EA42^/sov^2^* egg chambers versus N=0/47 controls). These findings are consistent with GSC hyperproliferation and/or failed differentiation. We also observed tumors and nurse cell nuclei residing within common follicles (Fig 3B) in about one-third of *sov* mutants (N=11/31 *sov^EA42^/sov^2^* versus N=0/24 in controls). This phenotype occurs when follicle cells either fail to separate egg chambers, or where those chambers fuse (Goode, Wright, and Mahowald 1992). Consistently, disorganization of the follicle cell monolayer was also observed (dashed lines, Fig 3B). In older *sov* ovaries, we also observed extensive cell death (Fig S1). Additionally, we found that rare *sov^EA42^/Y* males that escaped lethality had no germ cells (Fig S2). Thus, *sov* functions widely.

**Figure 3.**
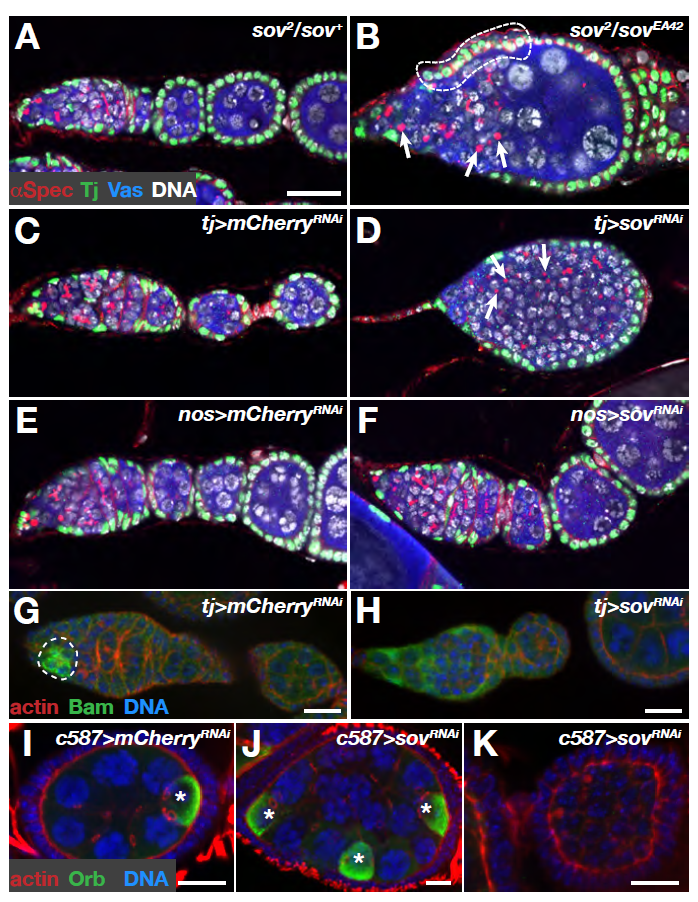
***sov*** **mutant defects.** Immunofluorescence for the indicated probes in the noted genotypes. (A–F) Single optical sections of germaria (4–5d posteclosion) stained with anti-Vas (blue), -Tj (green), -αSpectrin (red), and DAPI (white) with dot spectrosomes (arrows) and displaced follicle cells (dashed lines) shown. (G, H) Single optical sections of germaria (1d posteclosion) stained with anti-Bam (green), -actin (red), and DAPI (blue). Bam region of interest is shown (dashed circles). (I–K) Maximum intensity projections of germaria (1d posteclosion) stained with anti-Orb (green), -actin (red), and DAPI (blue). Orb+ cells shown (*). Bars: (A–F) 20μm, (G–K) 10μm.

To examine cell-type specific functions of *sov*, we used *tj-GAL4* to express *sov^RNAi^* (or *mCherry^RNAi^* controls) in somatic escort and follicle cells and *nanos-GAL4* (*nos-GAL4*) to express RNAi in the germline. Relative to the controls, the *tj>sov^RNAi^* germaria (Fig 3C, D) were abnormal, showing ovarian tumor phenotypes similar to those seen in *sov* mutants. Germaria were often filled with germline cells with dot spectrosomes and the follicle cells encroached anteriorly. Similar results were also observed using other somatic drivers (*c587-GAL4* and *da-GAL4*). Interestingly, *nos*>*sov^RNAi^* did not impair germarium morphology (Fig 3E, F) and egg chambers representing all 14 morphological stages of oogenesis appeared phenotypically normal. However, eggs produced from *nos>sov^RNAi^* mothers arrested during early embryogenesis, indicating that maternal *sov* is required for embryonic development. While *sov* expression patterns and germline RNAi suggest that *sov* is deployed in both the soma and germline, Wayne et al. (1995) reported *sov* to be somatic cell dependent. To further address a requirement for *sov* in the germline, we conducted germline clonal analysis. Depletion of *sov* from the germline results in an agametic phenotype (Fig S3). Additionally, *sov^RNAi^* resulted in embryonic germline defects in a high-throughput study (Jankovics et al. 2014). Taken together, these data indicate that there are both somatic and germline requirements for *sov* in oogenesis.

To explore the differentiation of the germline further, we examined the distribution of Bam, which is expressed in germarium region 1 germ cells (D. M. McKearin and Spradling 1990; D. McKearin and Ohlstein 1995; Ohlstein and McKearin 1997). In the absence of Bam, germ cells hyperproliferate, resulting in tumors composed of 2-cell cysts. Given the prevalence of undifferentiated tumorigenic germ cells upon somatic *sov^RNAi^* knockdown, we asked if *sov* has a nonautonomous role in promoting Bam expression in germ cells. Indeed, somatic knockdown of *sov* results in significantly less Bam expressed in germ cells relative to controls (Fig 3G, H; Fig S4). Both *tj-GAL4>sov^RNAi^* and *c587-GAL4>sov^RNAi^* resulted in a strong decrease in Bam expression. Germline depletion of *sov* did not alter Bam expression (Fig S4). The reduction in Bam expression in the germline is consistent with a nonautonomous role of Sov function in the soma for differentiation signals directed to the germline.

To examine the defective follicle encapsulation of the germline cysts following *sov* depletion, we used the oocyte-specific expression of Oo18 RNA binding protein (Orb) (Lantz et al. 1994) to count the number of oocytes per cyst. Orb specifies the future single oocyte at the posterior of the egg chamber in control egg chambers (Fig 3I, Fig S2D). Somatic depletion of *sov* resulted in examples of both egg chambers with either too many oocytes (Fig 3J) or no oocytes (Fig 3K, Fig S4). In a wild type 16-cell germline cyst, one of the two cells that has four ring canals becomes the oocyte, and this feature can be used to determine if extra germline divisions had occurred in cysts, or if multiple cysts were enveloped by the follicle cells. We saw that the egg chambers with multiple Orb+ cells always had more than 16 germ cells, and in the representative example shown, all three Orb+ cells had four ring canals, indicating that egg chamber fusion had occurred (Fig 3J). Taken together, we conclude that Sov functions in the soma to ensure proper differentiation of the germline.

### Sov represses gene expression in the ovary

To provide mechanistic insights into the function of Sov, we performed transcriptome profiling using triplicated Poly-A^+^ RNA-seq analyses of ovaries from *sov*, *sov*^RNAi^, and control females. The gene expression profiles of ovaries from sterile females were markedly different from controls, primarily due to derepression in the mutants (Fig 4A; Table S2). Among genes showing differences in expression (FDR *p*adj < 0.05), we found 1,752 genes with >4-fold increased expression in mutants, while there were only 172 genes with >4-fold decreased expression (Table S3). To explore where these derepressed genes are normally expressed, we examined their expression in other female tissues and in testes in a set of quadruplicated RNA-seq experiments (Fig 4B). Many of the genes suppressed by Sov in ovaries were highly expressed in other female tissues, but not testes, indicating that wild type Sov prevents ectopic gene expression.

**Figure 4.**
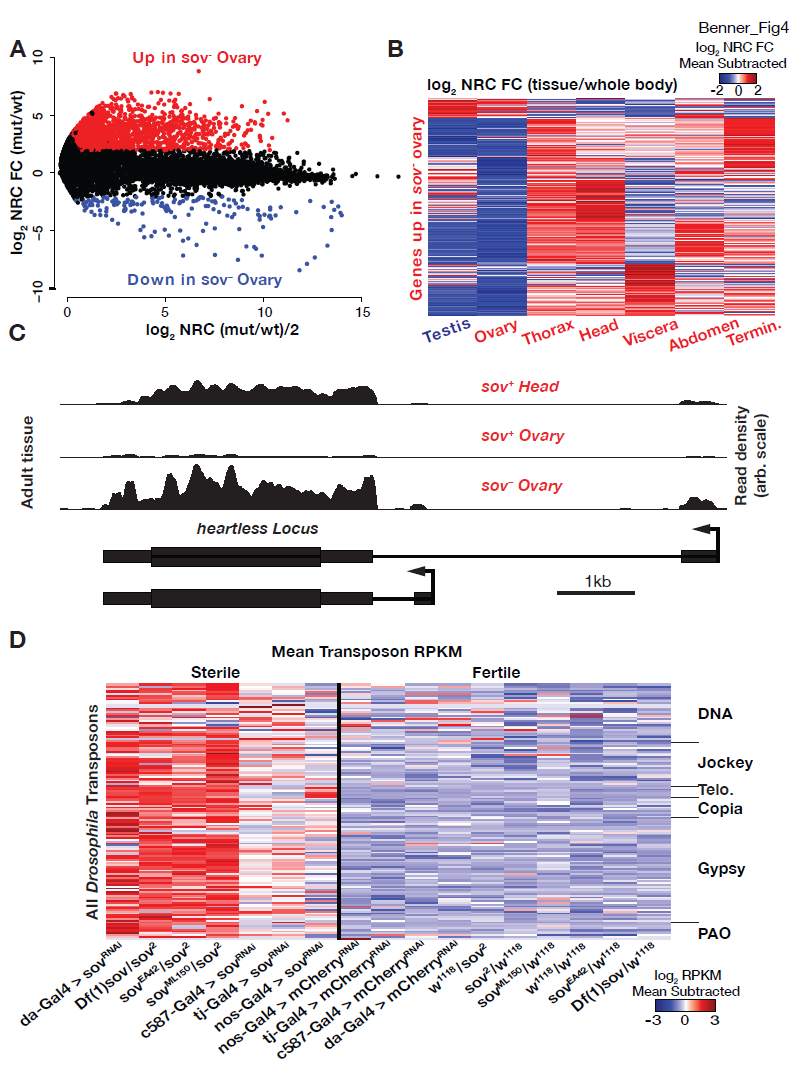
***sov*** **mutant transcriptome.** (A) Relative expression in *sov*^−^ vs *sov*^+^ ovaries plotted against the mean expression in both sample types. Units are log_2_ normalized read counts (NRC). Data points are genes, those with >4-fold change (log_2_ 2, red) or <–4 fold change (log_2_ −2, blue) and FDR *p*adj value <0.05 are highlighted. (B) Tissue-biased expression in wild type tissues for genes derepressed in *sov* mutant ovaries. Heatmap from mean-subtracted ratios scaled across tissues (red=higher; blue=lower). (C) RNA-seq normalized read densities of the *heartless* locus. (D) Transposable element expression in *sov* mutants (sterile) and controls (fertile). Heatmap from mean-subtracted reads (in RPKM. red=higher; blue=lower) scaled for each transposable element (rows) across genotypes (columns). Transposon classes for DNA, non-LTR (Jockey), telomeric repeat (Telo), and LTR (Gypsy, Copia, and PAO) are indicated.

To determine what types of genes are derepressed in *sov* mutants, we performed Gene Ontology (GO)(Gene Ontology Consortium 2015) term analysis (Fig S5; Table S4). The genes with lower expression in *sov* mutants had only a few significant GO terms that were predominantly oogenic in nature (as expected given the general lack of mature eggs). For example, there was poor expression of the chorion genes that are required to build the eggshell (Orr-Weaver 1991), as well as genes required for follicle cell development. In contrast, the derepressed genes in *sov* mutants showed a significant enrichment of cell signaling GO terms, including neuronal communication. We found that many genes repressed by Sov in the ovary, such as the *heartless* (*htl*) locus, which encodes a FGFR tyrosine kinase receptor important for neuron/glia communication (Stork et al. 2014), are indeed expressed in the head (Fig 4C). These results suggest that *sov* normally functions to repress a host of signaling pathways. Derepressed signaling in *sov* mutants is likely catastrophic for communication between various somatic cells and the germline during egg chamber development and could explain the variety of mutant ovarian phenotypes.

The general repressive role of *sov* was also revealed by a dramatic and coherent elevation of transposon expression in the mutants (Fig 4D; Table S3). Of the 138 transposable element classes detected in our gene expression profiling of *sov* mutants and controls, 91 had increased expression (FDR *p*adj < 0.05) >4-fold in *sov* mutants, while none had >4-fold decreased expression. The DNA, LINE, and LTR families of transposable elements were all derepressed in *sov* mutants. Interestingly, somatic knockdown of *sov* resulted in the most dramatic derepression of the *gypsy* and *copia* classes of transposons, which are *Drosophila* retroviruses that develop in somatic cells and are exported to the developing germline (Yoshioka et al. 1990). Thus, wild type *sov* may be important to protect the germline from infection. Equally interesting, germline knockdown resulted in the greatest derepression of the *HeT-A*, *TAHRE*, and *TART* transposons that compose the telomere (Mason, Frydrychova, and Biessmann 2008). This finding suggests that the general role of *sov* in silencing transposon expression includes control of both transposon functions necessary for normal cellular metabolism, exemplified by the telomere transposons, as well as detrimental activities of retrovirus-like elements. In addition to widespread deregulation of signaling and development pathways in *sov* mutants, transposon dysgenesis may contribute to the severe ovarian development defects in *sov* mutants.

### Sov is a suppressor of PEV and colocalizes with HP1

The repressive function of Sov is reminiscent of HP1a function as a general repressor of gene expression. HP1a was first characterized in *Drosophila* as a suppressor of PEV, a process that reduces gene expression due to spreading of heterochromatin into a gene region (Clark and Elgin 1992). To test the hypothesis that *sov* negatively regulates gene expression by promoting heterochromatin formation, we examined the role of *sov* in PEV in the eye, where patches of white^+^ (red pigmented) and white^−^ (not red pigmented) eye facets are easily observed (Fig 5A). If, like HP1a, Sov represses gene expression by promoting heterochromatin formation (Eissenberg et al. 1990), then it should suppress PEV, which is scored by increased eye pigmentation. We obtained five different variegating *w^+^* transgene insertions associated with either the heterochromatic pericentric region of chromosome arm 2L or spread along the length of the heterochromatin-rich chromosome 4. In control animals, these insertions show a characteristic eye variegation pattern (Fig 5B). Consistent with the allelic strengths seen in previous experiments, the weak *sov^2^* allele did not suppress PEV (Fig 5C), but the stronger *sov^ML150^*, *sov^EA42^*, and *Df(1)sov* mutations dominantly suppressed PEV (Fig 5D–F). These data demonstrate a role for Sov in heterochromatin formation.

**Figure 5.**
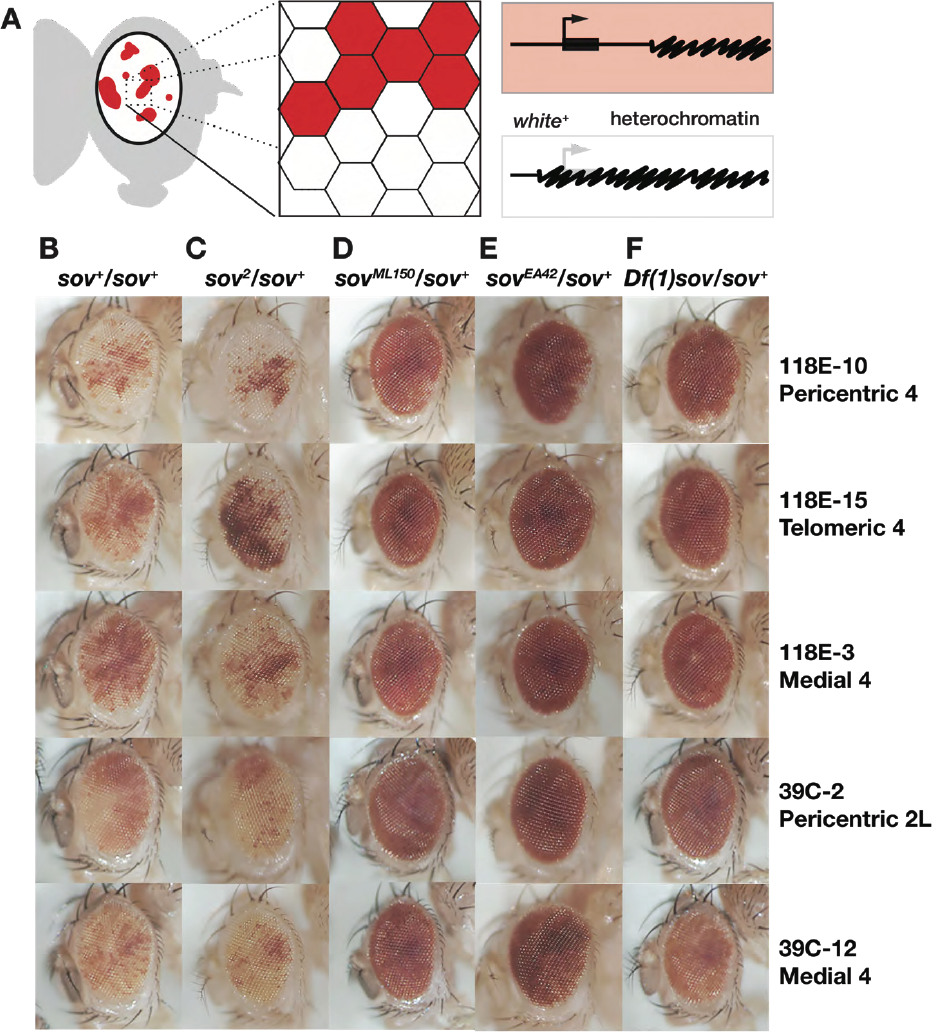
**Sov is a dominant suppressor of position effect variegation.** (A) Cartoon of position-effect variegation (PEV) in the eye. Expression of the *white* gene (bent arrow, thick bar, red) can be silenced (white) by proximal heterochromatin (squiggled) spreading. (B–F) Eyes from adults of the indicated genotypes (columns) with variegated expression of *P{hsp26-pt-T}* transgenes inserted into the indicated chromosomal positions (rows).

The strong repressive function of Sov is reminiscent of HP1a. Prior in vitro work suggests Sov may complex with HP1a (Alekseyenko et al. 2014). To determine if Sov and HP1a associate in vivo, we followed HP1a-RFP and GFP-Sov localization in 1–2 hr live embryos. During *Drosophila* embryogenesis, nuclei undergo rapid synchronous nuclear divisions prior to cellularization at cleavage division/nuclear cycle (NC) 14 (Foe and Alberts 1983). HP1a and Sov colocalized in all NC 10–12 embryos we examined (Fig 6A; N=7 embryos). During prophase, both HP1a and Sov were enriched in regions of condensed DNA (Fig 6A, 4:30). Whereas low levels of HP1a decorated DNA throughout division, Sov was depleted during mitosis (Fig 6A, 10:00 and 11:10). Upon reentry into interphase, Sov localization to nuclei resumed and was coincident with HP1a. Formation of heterochromatin is contemporaneous with or slightly proceeds HP1a apical subnuclear localization in NC 14 (Rudolph et al. 2007; Yuan and O’Farrell 2016). At this stage, we observed a strong colocalization of HP1a and Sov. Measuring the distribution of HP1a and Sov (Fig 6B, B’) confirmed high levels of colocalization within HP1a subnuclear domains (Fig 6C, shaded region). These data support the idea that HP1a and Sov are deposited maternally where they assemble into a complex.

**Figure 6.**
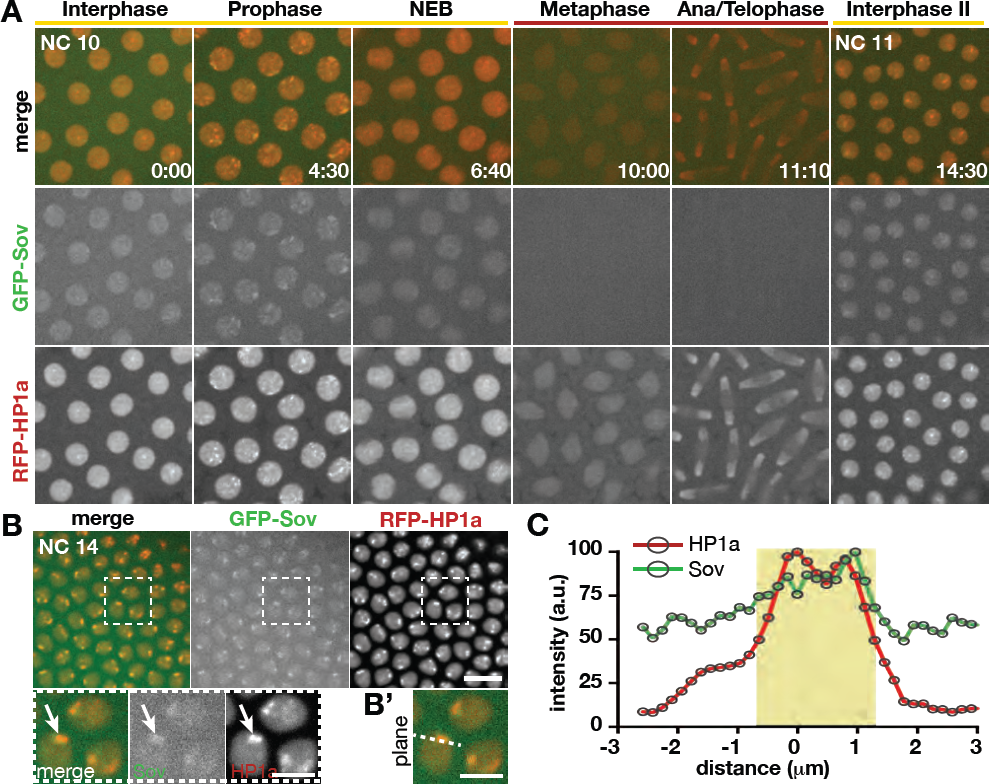
**Sov colocalizes with HP1a.** Stills from live imaging of embryos expressing GFPSov and RFP-HP1a. (A) Localization of GFP-Sov and RFP-HP1a (rows) in an embryo progressing from NC 10 to 11 with cell cycle stages (columns, nuclear envelope breakdown=NEB) and time (min:s) shown. (B) NC 14 embryo. Boxed regions are magnified in insets below. An HP1a subnuclear domain is shown (arrows). (B’) Single optical section containing peak HP1a fluorescence of inset from (B). Dashed line indicates region used for histogram analysis. (A,B) Maximum projections through 1.5 μm volume at 1F/30s. Bars: 10 μm; insets, 5 μm. (C) Histogram of HP1a and Sov fluorescence intensity measured in (B’). Fluorescence levels (arbitrary units) normalized to the peak fluorescence intensity for each channel and the distance (μm) to peak HP1a signal. Half maximum HP1a fluorescence is shaded (yellow).

## Discussion

### Sov is a novel heterochromatin-associated protein

Genomes contain large blocks of DNA of potentially mobile transposons and immobile mutated derivatives (Vermaak and Malik 2009). The cell keeps these transposons from wreaking havoc on the genome by actively suppressing their expression through condensation into heterochromatin. However, some tightly regulated expression from heterochromatin is required for normal cellular function. For example, the telomeres of *Drosophila* are maintained by transposition of mobile elements from heterochromatic sites (Mason, Frydrychova, and Biessmann 2008), and histone and rRNA genes are located within heterochromatic regions (Yasuhara and Wakimoto 2006). Weakening of heterochromatin by suppressors of variegation results in derepression of gene expression at the edges of heterochromatin blocks, suggesting that the boundaries between repressed and active chromatin can expand and contract (Weiler and Wakimoto 1995; Reuter and Spierer 1992). HP1a, encoded by *Su(var)*205, is a central component of heterochromatin (Ebert et al. 2006) that shows the same strong suppression of variegation that we observed in *sov/+* flies. Similarly, HP1a is also required to repress the expression of transposons (Vermaak and Malik 2009).

Repression is often stable and emerging themes suggest a robust set of activities, rather than a single component, maintain a chromatin state. For example, long-term repression is stabilized with complexes, such as the repressive Polycomb Group (PcG) which provides an epigenetic memory function (Kassis, Kennison, and Tamkun 2017). In order to create a stable epigenetic state, PcG complexes have subunits that modify histones and bind these modifications. Chromatin tethering of PcG complexes also involves DNA-binding proteins, such as the YY1-like ZnF protein Pleiohomeotic (Pho), which further reinforce localization. The DNA anchor proteins in PcG complexes are at least partially redundant with the histone-binding components (J. L. Brown et al. 2003), suggesting that localization is robust due to multiple independent localization mechanisms.

In mammals, HP1a also has both histone binding and DNA anchoring requirements. DNA anchoring is provided by a family of KRAB ZnF proteins that have undergone a massive radiation during evolution (Yang, Wang, and Macfarlan 2017; Ecco, Imbeault, and Trono 2017). KRAB proteins are essential for repression of transposons and more generally for a properly regulated genome. The KRAB family members use a KRAB-Associated Protein (KAP1) adapter protein to associate with HP1a and the SETDB1 methylase that modifies histones to enable HP1a binding (Fig 7). Although KRAB proteins have not been identified in *Drosophila*, the *Drosophila bonus* (*bon*) gene encodes a KAP1 homolog (Beckstead et al. 2005). *Drosophila* also has a SETDB1 encoded by *egg*, and loss of *egg* results in ovarian phenotypes reminiscent of those observed in *sov* loss-of-function females (Clough et al. 2007; Clough, Tedeschi, and Hazelrigg 2014; Wang et al. 2011). Like *sov* and *HP1a*, *bon* is a modifier of variegation, raising the possibility that they collectively coordinate gene regulation. In support of this hypothesis, Bon, Sov, and Egg were all identified in the same biochemical complex as HP1a (Alekseyenko et al. 2014). It is tempting to speculate that the single very long and ZnF-rich Sov protein plays the same direct binding role as the large family of KRAB ZnF proteins in mammals (Fig 7). If Sov uses subsets of fingers to bind DNA, it could localize to many different sequences in a combinatorial fashion. Additionally, the complex containing Sov and HP1a also contains RNA (Alekseyenko et al. 2014) and could act as a tether via a series of RNA intermediates as occurs in sex chromosome inactivation, for example (J. T. Lee 2009). As in the case of Pho in the PcG complexes, Sov might be a robustness factor rather than an absolute requirement, as we did not observe gross delocalization of HP1a following *sov^RNAi^* (not shown). Further work will be required to fully understand the relationship between Sov and HP1a.

**Figure 7.**
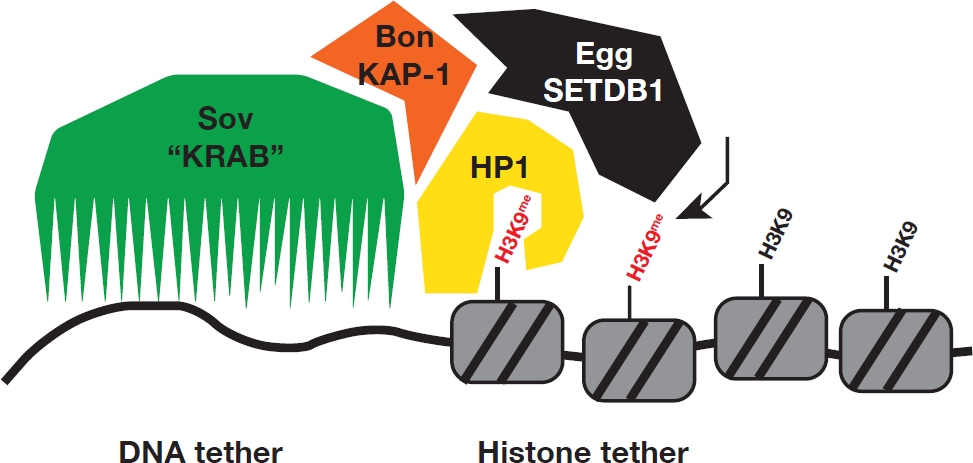
**Working model of** ***sov*** **function.** Sov protein function and attributes are analogous to the mammalian KRAB proteins (green). Like KRAB proteins, Sov is in complex with HP1a (yellow), Bon (mammalian KAP-1, orange) and Egg (mammalian SETDB1, black) providing a DNA tether to stabilize a repressed chromatin state.

### Repression by Sov promotes germline differentiation

The fact that there have been many female sterile alleles of *sov* suggests that the ovary is particularly sensitive to reduced *sov* activity. The female sterility phenotype is complex and somewhat variable, but partial *sov* loss of function is characterized by somatic-dependent differentiation defects. Stronger alleles also show a germline-dependent block in development and lethality. While *sov* functions seem diverse, the locus expresses a single major isoform, and Sov seems to be consistently located in the nucleus. We propose that the multifunctionality of Sov may be attributed to a single mechanism wherein Sov helps control heterochromatin formation required to repress ectopic gene expression and tissue-inappropriate responses in the ovary and repress transposons (Fig 7).

We observed dramatic derepression of transposons in *sov* mutants. Consistent with our observations, Czech et al (2013) reported a role of *sov* in transposon repression in a genomewide RNAi screen. That work raised the possibility that transposon repression by Sov occurs via the piRNA pathway (Brennecke et al. 2007; Yin and Lin 2007; Teixeira et al. 2017). However, genes involved more strictly with transposable element regulation, such as Piwi, do not have strong dosage effects on PEV (Gu and Elgin 2013), while *sov* and *HP1a*/*Su(var)205* do. This distinction suggests that Sov has a more general effect on heterochromatin, rather than specificity for transposon repression. Germline knockdown of HP1a results in a strong derepression of telomeric transposons (Teo et al. 2018), just like Sov. This is suggested to be due to a reduction in piRNAs specifically targeting these transposons and thus interplay between the piRNA pathway, Sov, and HP1a remains possible.

## Conclusions

Our data show that *sov* encodes a repressor of gene expression and transposons in the ovary. In the absence of *sov*, genes that are normally expressed at very low levels in the ovary, are activated. These same genes show dynamic patterns of expression in other adult tissues, suggesting that they are derepressed in the absence of Sov. The failure to restrict gene expression results in a range of phenotypes including ovarian tumors, defective oogenesis, and tissue degeneration. These data support the idea that Sov encodes a novel and essential protein that is generally repressive, likely via interactions with HP1a.

## Materials and Methods

We have adopted the FlyBase-recommended resources table which includes all genetic, biological, cell biology, genomics, manufactured reagents and algorithmic resources used in this study (Table S5).

### Flies and genetics

*sov* was defined by three X-linked, female sterile mutations including *sov^2^*, which mapped to ˜19 cM (Mohler 1977; Mohler and Carroll 1984), refined to 6BD (Wayne *et al*. 1995). We used existing and four custom-made deletions (*Df(1)BSC276*, *BSC285*, *BSC286* and *BSC297*) to map *sov* to the 4-gene *CG14438*–*shf* interval (Fig. 1; Table S5). *sov* mutations complemented *shf^2^*, but not *P{SUPor-P}CG14438^KG00226^* or *P{GawB}NP6070*, suggesting *CG14438* or *CR43496* is *sov*, which we confirmed by generating *Df(1)sov* (X:6756569.6756668;6770708) from FLP crossover between *P{XP}CG14438^d07849^* and *PBac{RB}e03842 (Parks et al. 2004; Cook et al. 2012)* to remove only those two genes.

We used the dominant female sterile technique for germline clones (Chou and Perrimon 1996). Test chromosomes, that were free of linked lethal mutations by male viability (*sov^2^)* or rescue by *Dp(1;3)DC486* (*sov^EA42^* and *Df(1)sov*), were recombined with *P{ry^+t7.2^=neoFRT}19A*, and verified by complementation tests and PCR. We confirmed *P{ry^+t7.2^=neoFRT}19A* functionality by crossing to *P{w^+mC^=GMR-hid}SS1, y^1^ w^*^ P{ry^+t7.2^=neoFRT}19A; P{w^+m*^=GAL4-ey.H}SS5, P{w^+mC^=UAS-FLP.D}JD2* and scoring for large eye size. We crossed females with FRT chromosomes to *P{w^+mC^=ovoD1-18}P4.1, P{ry^+t7.2^=hsFLP}12, y^1^ w^1118^ sn^3^ P{ry^+t7.2^=neoFRT}19A/Y* males for 24 hours of egg laying at 25°C. We heat-shocked for 1hr at 37°C on days 2 and 3. We dissected females (5d posteclosion) to score for *ovo^D1^* or wild type morphology.

We generated *PBac*{*GFP-sov}* from P[acman] BAC clone CH322-191E24 (X:6753282– 6773405) (Venken et al. 2009) grown in the SW102 strain (Warming et al. 2005). In step one, we integrated the positive/negative marker CP6-RpsL/Kan (CP6 promoter with a bi-cistronic cassette encoding the RpsL followed by the Kan), PCR-amplified with primers N-CG14438-CP6-RN-F and -R between the first two codons of sov, and selected (15 μg/ml Kanamycin) We integrated at the galK operon in DH10B bacteria using mini-lambda-mediated recombineering (Court et al. 2003). We amplified DH10B::CP6-RpsL/Kan DNA using primers N-CG14438-CP6-RN-F and -R. Correct events were identified by PCR, as well as resistance (15 µg/ml Kanamycin) and sensitivity (250 µg/ml Strep). In step two, we replaced the selection markers with a multi-tag sequence (Venken et al. 2011), tailored for N-terminal tagging (N-tag) and counter-selected (250 µg/ml Strep). The N-tag (3xFlag tag, TEV protease site, StrepII tag, superfolder GFP, FlAsH tetracysteine tag, and flexible 4xGlyGlySer (GGS) linker) was *Drosophila* codon optimized in a R6Kγ plasmid. We transformed plasmid into EPI300 for copy number amplification. We confirmed correct events by PCR and Sanger DNA sequencing. Tagged P[acman] BAC clone DNA was injected into *y^1^ M{vas-int.Dm}ZH-2A w*; PBac{y+-attP-3B}VK00033* embryos, resulting in *w^1118^; PBac{y[+mDint2] w^+mC^=FTSF.GFP-sov}VK00033*.

### Microscopy

We fixed ovaries in 4 or 5% EM-grade paraformaldehyde in PBS containing 0.1 or 0.3% Triton X-100 (PBTX) for 10–15min, washed 3x 15 min in PBTX, and blocked >30min in 2% normal goat serum, 0.5-1% bovine serum albumin (BSA) in PBS with 0.1% Tween-20 or 0.1% Triton X-100. Antibodies and DAPI were diluted into blocking buffer. We incubated in primary antibodies overnight at 4°C and secondaries 2–3hr at room temperature. Embryos (1–2 hr) were prepared for live imaging in halocarbon oil (Lerit et al. 2015). We imaged ovaries and embryos using a Nikon Ti-E system or Zeiss LSM 780 microscope and eyes with a Nikon SMZ.

### DNA-Seq

Genomic DNA was extracted from 30 whole flies per genotype (Huang, Rehm, and Rubin 2009)(Sambrook and Russell 2006) to prepare DNA-seq libraries (Nextera DNA Library Preparation Kit). We used 50 bp, single-end sequencing (Illumina HiSeq 2500, CASAVA base calling). Sequence data are available at the SRA (SRP14438). We mapped DNAseq reads to FlyBase r6.16 genome with Hisat2 (-k 1–no-spliced-alignment)(Kim, Langmead, and Salzberg 2015). We used mpileup and bcftools commands from SAMtools within the genomic region X:6756000–6771000 (H. Li et al. 2009; H. Li 2011) for variant calling and snpEFF to determine the nature of variants in *sov* mutants (Cingolani et al. 2012).

### RNA-seq

Stranded PolyA+ RNA-seq libraries from *sov* and control ovaries (Table S5) were created (H. Lee, Cho, et al. 2016) and are available at GEO (GSE113977). We extracted total RNA (Qiagen RNeasy Mini Kit) in biological triplicate from 15 ovaries (4–5d posteclosion) and used 200 ng with 10 pg of ERCC spike-in control RNAs (pools 78A or 78B) for libraries (Jiang et al. 2011; Zook et al. 2012; Pine et al. 2016; H. Lee, Pine, et al. 2016). We used 50 bp, single-end sequencing as above. Tissue expression analysis are from GEO accession GSE99574, a resource for comparing gene expression patterns.

We mapped RNA-seq reads to FlyBase r6.21 with Hisat2 (-k 1–rna-strandness R–dta) (Kim, Langmead, and Salzberg 2015). We determined read counts for each attribute of the FlyBase r6.21 GTF file (with ERCC and transposable element sequences), with HTSeq-count (Anders, Pyl, and Huber 2015). Transposon sequences were from the UCSC Genome Browser RepeatMasker track (Casper et al. 2018; Smit, Hubley, and Green 2013-2015).

We conducted differential expression analysis with DESeq2 (pAdjustMethod = “fdr”) (Love, Huber, and Anders 2014). We removed genes with read counts <1 and read counts for transposable elements with >1 locations were summed for the DESeq2 analysis. *Df(1)sov/sov^2^* replicate 3 and *sov^2^/w^1118^* replicate 1 were failed. For *sov* mutant *vs.* control DESeq2 analysis, all *sov* mutants were compared to all wild type controls. For tissue types, we compared each sexed tissue to sexed whole organism. We used reads per kb per million reads (RPKM) for gene-level expression.

For heatmaps, we calculated Euclidean distance, performed hierarchical cluster analysis (agglomeration method = Ward), and mean-subtracted scaled across genotypes. Mean sample RPKM correlation values (Table S1) were calculated by cor.test() function with Pearson correlation coefficient in R (R Core Team 2017). We represented derepressed genes as mean-subtracted scaled values across tissues in the heatmap (Table S6).

For read density tracks, replicate raw read files were combined. Bedgraph files were created with bedtools genomecov (Quinlan and Hall 2010) visualized on the UCSC genome browser (Kent et al. 2002). Tracks were scaled by the number of reads divided by total reads per million.

We used ClueGO (Bindea et al. 2009), with Cytoscape (Shannon et al. 2003) for one-sided enrichment analysis (see Table S4).

Images were assembled using ImageJ and Photoshop. We used ROI tool in ImageJ, plotted/analyzed image data (Microsoft Excel and GraphPad Prism), and calculated significance by D’Agnostino and Pearson normality tests, followed Student’s two-tailed t-test or Mann-Whitney tests.

## Acknowledgements

Stocks obtained from the Bloomington Drosophila Stock Center (NIH P40OD018537) and Drosophila Genomics and Genetic Resources at the Kyoto Institute of Technology were used in this study. Monoclonal antibodies were obtained from the Developmental Studies Hybridoma Bank, created by the NICHD of the NIH and maintained at The University of Iowa, Department of Biology, Iowa City, IA 52242. Sequencing was performed by the NIDDK Genomics Core, under the direction of Harold Smith. Genetic and genomic information was obtained from FlyBase. This work utilized the computational resources of the NIH HPC Biowulf cluster (http://hpc.nih.gov). We thank BACPAC resources for plasmids, Norbert Perrimon for the *sov^EA42^* stock and Don Court, Neal Copeland, and Nancy Jenkins for the recombineering strain. We thank Astrid Haase, Nasser Rusan, David Katz, and members of the Oliver and Lerit labs for stimulating discussions. Particular thanks go to Stacy Christensen and Kimberley Cook for generating and characterizing the *sov* deletions. This research was supported in part by the Intramural Research Program of the NIH, National Institute of Diabetes and Digestive and Kidney Diseases (NIDDK) to BO. DAL and EAC were supported by NIH grant 5K22HL126922-02 (DAL). LB was supported by the NIH Graduate Partners Program. KJTV was supported by Baylor College of Medicine, the Albert and Margaret Alkek Foundation, the McNair Medical Institute at The Robert and Janice McNair Foundation, the March of Dimes Foundation (#1-FY14-315), the Foundation for Angelman Syndrome Therapeutics (FT2016-002), the Cancer Prevention and Research Institute of Texas (R1313), and NIH grants (1R21HG006726, 1R21GM110190, 1R21OD022981, and R01GM109938). CW and KRC were supported by the Office of the NIH Director, the National Institute of General Medical Sciences and the National Institute of Child Health and Human Development (P40OD018537). KRC was supported by the National Center for Resource Resources (R24RR014106) and the National Science Foundation (DBI-9816125).

## Author contributions

LB, KC, BO, and DAL designed the project and analyzed data. LB, KC, CW, and KJTV performed genetics. LB and HY performed genomics. LB, EAC, CW and DAL performed imaging. LB, BO, and DAL wrote the manuscript. All authors reviewed data and provided feedback on the manuscript.

